# Do cooperatively breeding mammals live longer? A re-appraisal

**DOI:** 10.1101/769455

**Authors:** Jack Thorley

## Abstract

Recent comparative studies have suggested that cooperative breeding is associated with increases in maximum lifespan among mammals, replicating a pattern also seen in birds and insects. In this study, I re-examine the case for increased lifespan in mammalian cooperative breeders by analysing a large dataset of maximum longevity records. Unlike one previous study, I found no consistent, strong evidence that cooperative breeders have longer lifespans than other mammals, after having controlled for variation in body mass, mode of life and data quality. The only exception to this general trend was displayed by the African mole-rats (the Bathyergid family): all members of this family are relatively long-lived, but the social, cooperatively breeding species appear to be much longer-lived than the solitary species, the latter having not been known to live beyond 11 years in captivity. However, solitary mole-rat species have rarely been kept in captivity or followed longitudinally in the wild, and so it seems likely that their maximum lifespan has been grossly underestimated when compared to the highly researched social species. As few other subterranean species have received much attention in a captive or wild setting, I also suggest that current data also makes it impossible to rule out a causal role of subterranean living on lifespan extension in mammals, and that any future studies wanting to test for such an association should wait until more high quality longevity data is available from a wider range of permanently subterranean species.

## INTRODUCTION

Comparative studies of birds and insects have shown that cooperative and eusocial breeding systems are associated with extended lifespans (Arnold & Owens, 1998; Beauchamp, 2014; Downing *et al.*, 2015; Keller & Genoud, 1997). A common explanation for the association between these breeding systems and increased lifespan is that group-living is associated with a reduction in extrinsic mortality in breeding individuals, which selects for greater longevity (Lucas and Keller 2019). This is thought to be the case in eusocial insect societies, with the ‘queens’ of many ant, bee and termite species living in a sheltered nest that is defended against predators by a large workforce (Carey 2001; Hölldobler & Wilson 1990). An alternative possibility is that high annual survival increases local breeding competition, leading to delayed dispersal, family-living, and helping (Brown, 1987; Griesser *et al.*, 2017), such that having a relatively long lifespan makes cooperative breeding more likely to evolve, and phylogenetic reconstructions of cooperative breeding supports this argument in birds (Downing *et al.*, 2015).

Observations from certain mammals are also suggestive of a relationship between cooperative breeding and lifespan. The social mole-rats (family Bathyergidae), for example, include some of the longest-lived mammals for their size, with the naked mole-rat *Heterocephalus glaber*, a 40g species from the Horn of Africa providing the most extreme case. In this species, breeding females can live for more than three decades in captivity and show no apparent age-related changes in physiology or mortality rate (Buffenstein, 2005; Fang *et al.*, 2014; Kim *et al.*, 2011; Ruby *et al.*, 2018; but see Dammann *et al.* 2019), prompting much interest in naked mole-rats as a model organism for gerontological research. However, as well as being long-lived, mole-rats are completely subterranean and are thought to seldom be exposed to predators in their closed-off burrow systems (Dammann & Burda 2007; Sándor 2017), raising the question of whether it is their subterranean lifestyle, their cooperative breeding, or both that contribute to their longevity. To try and separate the role of subterranean living and cooperative breeding on maximum lifespan, a recent comparative analysis (Healy, 2015) examined the role of cooperative breeding and fossoriality (defined as species which make extensive use of burrow systems, but forage either above or below ground) on lifespan across terrestrial mammals and found that, when both traits were considered together in a phylogenetically controlled analysis, only cooperative breeding was significantly associated with longer lifespan.

While the above study documented a positive relationship between mammalian cooperative breeding and maximum lifespan, others have failed to find a relationship. Using phylogenetically independent contrasts, Lukas and Clutton-Brock (2012a) compared cooperatively breeding mammals to socially monogamous mammals and found no clear difference in maximum lifespan between species with these two mating systems. Since cooperative breeding is sparsely represented across the mammalian clade (less than 1% of mammals, Lukas & Clutton-Brock, 2012b), slight differences in data availability or analytical method or contrasts in definitions could contribute inconsistencies in results.

In this study, I re-analysed the variation in maximum lifespan in terrestrial mammals with an extended dataset to re-assess whether cooperative breeding is associated with enhanced longevity. Although the use of maximum lifespan records as a proxy for lifespan is not without criticism (e.g. Baylis et al. 2014), the absence of detailed life table information for many species usually precludes the use of alternative ageing metrics in a comparative setting. My approach deviates from the previous study of this topic by Healy (2015) in several important respects. The first major difference is in the categorisation of species according to their use of the underground environment. Here, I separate species according to whether they are subterranean feeders or not. Subterranean feeders spend almost their entire lives underground and are therefore likely to experience the low levels of predation and extrinsic mortality that should select for longer lifespans (Hartman, 1995; Novikov & Burda 2013). In contrast, Healy chose a more relaxed definition whereby all mammals that make extensive use of the underground environment were classed as fossorial. This definition forces one to group subterranean feeders (such as the moles, mole-rats, pocket gophers, or coruro *Spalacopus cyanus*) with species that use the underground environment extensively for resting, denning, and sometimes food storage, but which otherwise spend a large part of their daily activity period foraging above-ground (for example armadillos, aardvark *Orcyteropus afer*, or Eurasian badger *Meles meles*). It is not clear that mammals falling into the latter category should experience reduced predation rates to the same extent as subterranean feeders (see Healy *et al.*, 2014), and grouping the two classes could therefore prevent the detection of an effect of subterranean living on lifespan. Secondly, the current study controls for sample size (as others have done: Kamilar *et al.*, 2010; Minias & Podlaszczuk, 2017), a factor not included in Healy’s analysis. Since cooperative breeders are more frequently studied and larger numbers of individuals have been kept in captivity, maximum lifespan estimates for cooperative breeders are likely to be greater than for non-cooperatively breeding species for purely numerical reasons (see methods) – so sample size needs to be controlled for (Moorad *et al.*, 2012). Thirdly, by focussing on ground-dwelling (i.e. non-arboreal) mammals, Healy disregards lifespan data from the callitrichid primates (the marmosets and tamarins), which represent a sizeable fraction of the total pool of cooperatively breeding species. Here, I include information from callitrichid primates.

This study consisted of two stages. The first stage carried out a large-scale analysis of maximum lifespan across 719 mammals, controlling for subterranean living, data quality, and body mass. The application of different phylogenetic modelling techniques to this dataset suggested that the distribution of data across categorical predictors might have affected the ability of the models to discriminate between the effect of cooperative breeding and subterranean living on lifespan. To address this concern, lifespan variation was also analysed at the taxonomic level of the family, since five of the mammalian families that contain cooperatively breeding species also contain non-cooperatively breeding species with which they could be compared (the bathyergid mole-rats, the callitrichid primates, the canids, the cricetid rodents, and the herpestid mongooses). If cooperative breeding is consistently associated with increased lifespan, a relationship between cooperative breeding and lifespan should be present in each family.

## METHODS

### Global dataset

Maximum lifespan data was taken from the AnAGE database for 719 terrestrial mammal species (Figure 1; De Magalhães & Costa, 2009). This dataset provides estimates of the sample size for each longevity record, reflecting orders of magnitude in the number of specimens that contributed to the record. Species with fewer than 10 specimens contributing to their record (‘tiny’) were excluded. The only exception to this data restriction was the Cape mole-rat *Georychus capensis*, which was retained because of the general lack of longevity data for African mole-rats, family Bathyergidae; for model fitting purposes this species was reclassified as having ‘small’ sample size. This left 323 species with a ‘small’ sample size (10-100 specimens), 272 species with a medium sample size (100-1000), and 123 species with a ‘large sample size’ (over 1000 specimens). Sampling was not evenly distributed across cooperative and non-cooperative species: for cooperative species, large, medium and small lifespan sample sizes reflected 39.3%, 39.3% and 21.4% of species, as compared to 16.2%, 37.8% and 46.0% for non-cooperative species (χ^2^_2_ = 12.01, p = 0.002). All else being equal then, longevity records for cooperative breeders are likely to be higher simply due to sampling effort. Cooperatively breeding species were defined as those species where a proportion of females do not breed regularly and have been shown to perform alloparental care in the form of offspring provisioning (as per Lukas & Clutton-Brock, 2012a, 2017; Solomon & French, 1997; n = 28 cooperative breeders). For each species further information was added on adult body mass, subterranean living and habitat. Subterranean living demarcated species as subterranean or non-subterranean feeders using information from several sources (Begall *et al.*, 2007; Mittermeier *et al.*, 2018; n = 13 subterranean species). For ‘habitat’, species were defined as arboreal (n = 150), semi-arboreal (n = 70) or terrestrial (n = 499), using information from *Walker’s Mammals of the World* (Nowak, 1999) and the *Handbook of the Mammals of the World* (Mittermeier *et al.*, 2018). The updated mammalian phylogeny from Fritz *et al.* (2009) was used across all analyses.

**Figure 1.**
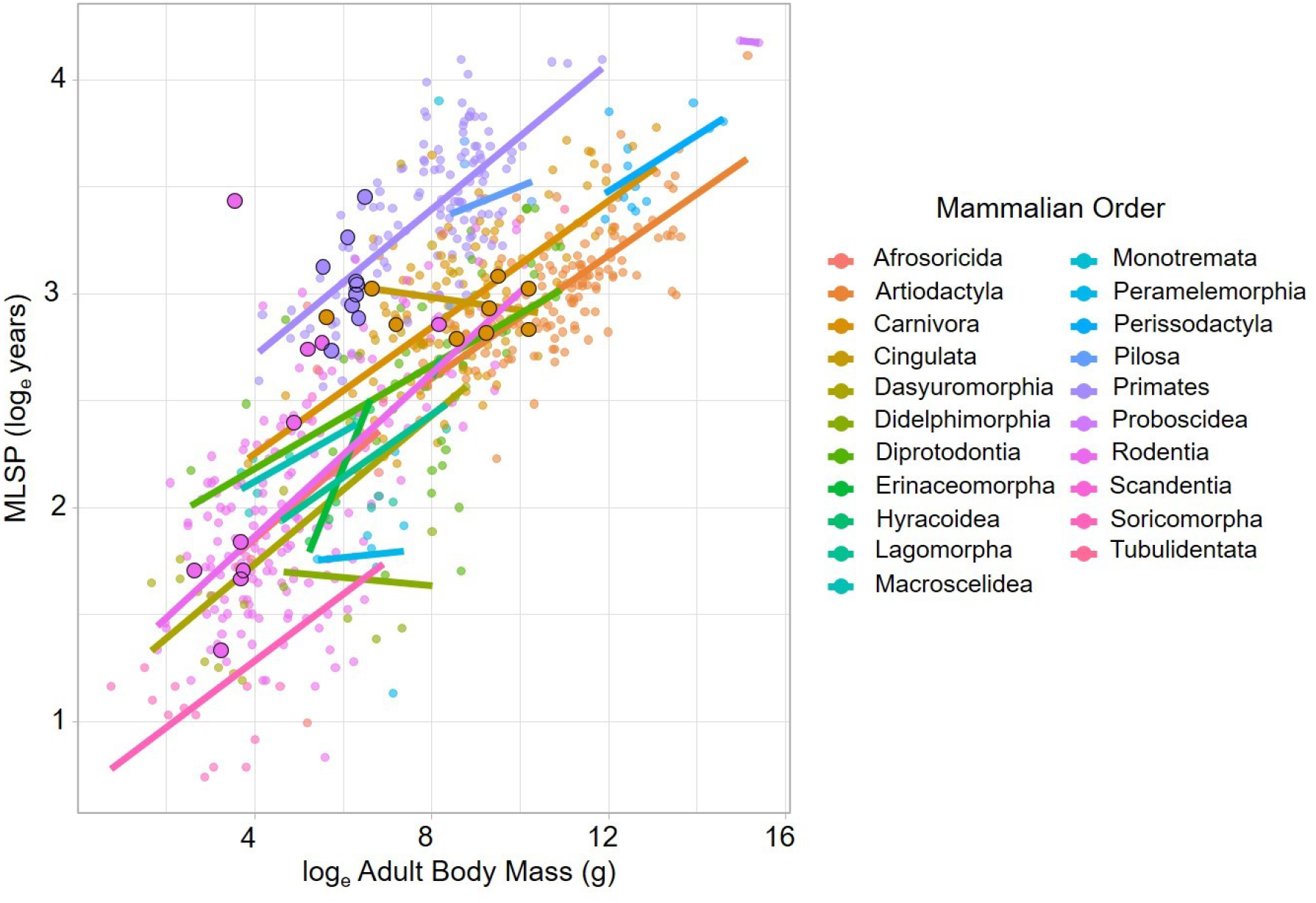
The relationship between loge adult body mass and loge maximum lifespan for 719 terrestrial mammals. Species are coloured according to Order, and cooperative breeders have been given larger, highlighted points. An ordinary least-squares regression is fitted through each Order to indicate phylogenetic differences in the scaling relationship.

### Global analysis

To investigate the influence of the chosen predictors on maximum lifespan, a global model was fitted with two different comparative methods.

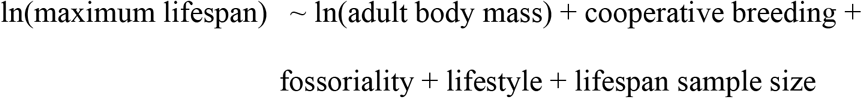

Adult body mass was z-score transformed before model fitting. Method 1 took a phylogenetic generalised least squares (PGLS) approach using the R package *phylolm* (Tung Ho & Ané, 2014). PGLS is a modification of generalised least squares method, which incorporates information from a phylogeny to generate parameter estimates that account for the expected covariance of traits that is due to shared ancestry. The phylogenetic nonindependence is calculated through lambda (λ), a multiplier of the off-diagonal elements of the expected variance-covariance matrix. When λ = 0, the off-diagonal elements are set to zero and phylogenetic dependency is completely absent, whereas λ = 1 implies a strong phylogenetic signal in the data that is structured according to a Brownian motion model of trait evolution. A PGLS model was also fitted in the *caper* package (Orme *et al.*, 2018) to act as an additional comparison. Method 2 implemented a Bayesian phylogenetic mixed model (PLMM) approach using the R package *MCMCglmm* (Hadfield, 2010). PLMMs account for nonindependence between species by incorporating the phylogenetic tree as a random effect (Hadfield & Nakagawa, 2010). The model was run for 115000 iterations with a burn-in of 15000 and a thinning interval of 100. An inverse Wishart prior was chosen for the variance components (V = 1, nu = 0.02). Diagnostics confirmed an adequate mixing of chains for these settings.

### Family-level analysis

Maximum lifespan variation was analysed at the level of the family for five mammalian families containing both cooperatively breeding and non-cooperatively breeding species: the bathyergid mole-rats, the callitrichid primates, the canids, the cricetid rodents, and the herpestid mongooses. For the Callitrichidae, Canidae, Cricetidae, and Herpestidae, a general linear model was fitted of the form: ln(maximum lifespan) ~ ln(adult body mass) + lifespan sample size. For the Bathyergidae, the lifespan sample size was excluded as only six species were present in the dataset. The residuals from each analysis represent the residual lifespan after controlling for body mass and sample size. Emphasis is placed on visualisation of the residuals for the Bathyergidae and the Herpestidae for sample size reasons. For the remaining three families, t-tests were carried out on the residuals to compare cooperatively breeding and non-cooperatively breeding species.

## RESULTS & DISCUSSION

Across all phylogenetic models fitted to the global dataset, cooperative breeding was positively associated with maximum lifespan (Table 1), but in all cases the effect size was small, reflecting at most a 2.8% increase in lifespan, and in two cases, the term was non-significant (*pgls*, p = 0.091; MCMC*glmm*, p = 0.089, phylolm). In contrast, subterranean living was associated with an 8.9% increase in maximum lifespan (*pgls*, p = 0.019; *MCMCglmm*: p = 0.015), though with one approach, the term was non-significant (*phylolm*, p = 0.096). That the three models disagreed in their attribution of significance to model terms despite similar effect sizes suggests that it was difficult for the models to apportion variance in lifespan to either cooperative breeding or subterranean living. It is likely that this is an artefact of both traits being rarely represented in the dataset, and when present, often co-occurring in the same taxa (Mundry, 2014).

**Table 1.** Global analyses of maximum lifespan across terrestrial mammals. Models were fitted using two phylogenetic methods. Significant model terms are highlighted in bold. For MCMCglmm, terms with credible intervals which do not overlap zero are deemed biologically significant.

| Model Term | PGLS, caper |  | PGLS, phylolm |  | PGLMM, MCMCglmm |
| --- | --- | --- | --- | --- | --- |
|  | Estimate (SE) | p-value | Estimate (SE) | p-value | Estimate (95% CI) |
| Intercept | 3.098 (0.327) |  | 3.117 (0.459) |  | 3.113 (2.49 – 3.75) |
| Adult Body Mass | **0.314 (0.026)** | **<0.001** | **0.249 (0.029)** | **<0.001** | **0.316 (0.27 – 0.36)** |
| Cooperative Breeder: non-cooperative | -0.069 (0.041) | 0.091 | **-0.088 (0.034)** | **0.011** | -0.069 (-0.14 – 0.01) |
| Fossoriality: non-subterranean | **-0.299 (0.127)** | **0.019** | -0.279 (0.169) | 0.096 | **-0.301 (-0.55 – - 0.06)** |
| Lifestyle: semi-arboreal | 0.081 (0.045) | 0.072 | 0.071 (0.054) | 0.119 | 0.082 (0.00 – 0.17) |
| Lifestyle: arboreal | **0.121 (0.051)** | **0.019** | 0.105 (0.062) | 0.089 | **0.124 (0.02 – 0.22)** |
| Sample size: medium | **0.125 (0.019)** | **<0.001** | **0.092 (0.017)** | **<0.001** | **0.124 (0.09 – 0.16)** |
| Sample size: large | **0.205 (0.025)** | **<0.001** | **0.183 (0.024)** | **<0.001** | **0.205 (0.16 – 0.26)** |
| | $\lambda = 0.971 (0.965-0.981)$ | | $\sigma^2 = 0.0044$ | | Residual var = 0.012<br>(0.009 – 0.016, 95% CI)<br>Phylogenetic variance = 0.378<br>(0.327 – 0.440, 95% CI) |

The lack of a clear relationship between cooperative breeding and maximum lifespan was further supported by analyses conducted on specific mammalian families (Figure 2). After correcting for body mass and sample size, residual maximum lifespan did not vary with cooperative breeding in the callitrichid primates (Welch’s t-test, t = 0.22, p = 0.83), the canids (t-test, t = 1.26, p = 0.24), the cricetid rodents (t-test, t =−0.46, p = 0.66), or the herpestid mongooses (visual inspection of Figure 2). A consideration of the social organisation of cooperative societies can make sense of this null result. Cooperative mammal societies are characterised by high rates of reproduction (in many species, dominant females produce relatively large litters and breed several times per year) and intense intrasexual competition (Barrette *et al.*, 2012; Clutton-Brock, 2016; Clutton-Brock *et al.*, 2006), both features which would classically be expected to reduced lifespan (Stearns, 1989). In addition, a previous study found no link between group size and lifespan in mammals (Kamilar *et al.*, 2010), so it is not clear that the large group size of cooperative breeders necessarily contributes to their pace of life either.

**Figure 2.**
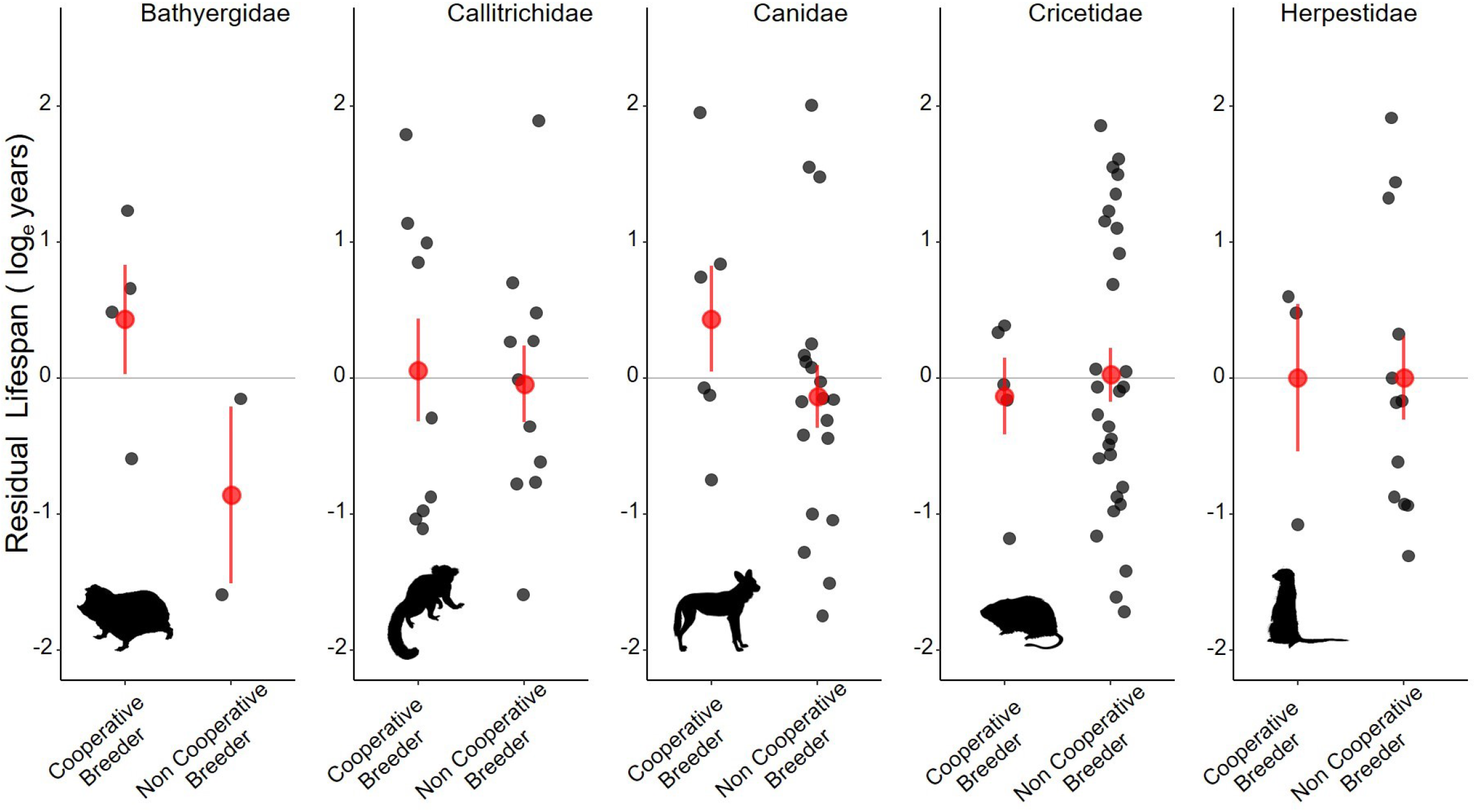
The residual lifespan of cooperative breeders compared to non-cooperative breeders in the families containing several cooperative and non-cooperative breeders. Residuals represent the remaining variation in maximum lifespan after the effects of body mass and lifespan sample size have been accounted for in a general linear model. (The Bathyergidae model only included body mass as a predictor). Black points represent raw standardised residuals, while red points detail the mean ± 1 sem. All pictures taken from Phylopic (http://phylopic.org) are free to use under Creative Commons license.

The only family where cooperatively breeding species appear to live longer is in the bathyergid mole-rats. Within this family the social taxa such as the naked mole-rat (31-year maximum lifespan) and Damaraland mole-rat *Fukomys damarensis*, (15.5 years) live markedly longer than the non-social, solitary members of the family such as the Silvery mole-rat *Heliophobius argentocinereus*, (7.5 years) or the Cape mole-rat (11.2 years). However, this comparison must be treated with caution, for unlike the social species, solitary mole-rats have attracted much less interest from researchers and are notoriously difficult to maintain in captivity, which will lead to large underestimates of the longevity of solitary species.

The results from the global analyses are more suggestive of a role of subterranean living on lifespan, but again, few subterranean mammals have been kept in captivity or been the focus of long-term individual-based studies. It therefore seems premature to place judgement on the role of subterranean living on ageing patterns until higher resolution data is collated from a larger number of species which permanently inhabit a subterranean niche, both in the wild and in captivity. This being the case, it not it is not particularly useful, for example, to claim that placental moles are on average characteristically short-lived in the context of all mammals (as per Williams & Shattuck 2015). For one, moles are rarely studied, and a cursory glance at another ageing database (Jones *et al.* 2009) indicates that one of the few well-studied species, the European mole *Talpa europaea* (not present in this study), can live for seven years, a relatively long time for an 80g insectivore. There are also other illustrative cases outside of the African mole-rats that indicate a possible association between subterranean living and lifespan extension. For example, the subterranean and solitary plains pocket gopher (*Geomys bursarius*) and the Middle East blind mole rat (*Nannospalax ehrenbergi*) have been known to live for 12 years and 15 years respectively (Weigl 2005).

Future studies assessing the role of sociality or mode of life on lifespan variation in should consider the availability and quality of data before conducting large-scale comparative analyses. Where data is scant, or when traits are unevenly distributed across the tree of life, researchers might often be better served conducting more focused analyses that compare closely related species whose ecology differs little beyond the trait of interest; as carried out in this study in the context of mammalian cooperative breeding.

## ACKNOWLEDGEMENTS

JT would like to thank Prof. Tim Clutton-Brock for his comments on the manuscript. The work was carried out with the support of a Natural Environment Research Council Doctoral Training Program. There are no conflicts of interest related to the work

